# Mechanochemical coupling and junctional forces during collective cell migration

**DOI:** 10.1101/558452

**Authors:** J. Bui, D. E. Conway, R. L. Heise, S.H. Weinberg

## Abstract

Cell migration, a fundamental physiological process in which cells sense and move through their surrounding physical environment, plays a critical role in development and tissue formation, as well as pathological processes, such as cancer metastasis and wound healing. During cell migration, dynamics are governed by the bidirectional interplay between cell-generated mechanical forces and the activity of Rho GTPases, a family of small GTP-binding proteins that regulate actin cytoskeleton assembly and cellular contractility. These interactions are inherently more complex during the collective migration of mechanically coupled cells, due to the additional regulation of cell-cell junctional forces. In this study, we present a minimal modeling framework to simulate the interactions between mechanochemical signaling in individual cells and interactions with cell-cell junctional forces during collective cell migration. We find that migration of individual cells depends on the feedback between mechanical tension and Rho GTPase activity in a biphasic manner. During collective cell migration, waves of Rho GTPase activity mediate mechanical contraction/extension and thus synchronization throughout the tissue. Further, cell-cell junctional forces exhibit distinct spatial patterns during collective cell migration, with larger forces near the leading edge. Larger junctional force magnitudes are associated with faster collective cell migration and larger tissue size. Simulations of heterogeneous tissue migration exhibit a complex dependence on the properties of both leading and trailing cells. Computational predictions demonstrate that collective cell migration depends on both the emergent dynamics and interactions between cellular-level Rho GTPase activity and contractility, and multicellular-level junctional forces.

## INTRODUCTION

Cell migration is a fundamental physiological process in which cells sense and move through their surrounding physical environment. Adherent cell migration plays a critical role in development and tissue formation, as well as pathological processes, such as cancer metastasis and wound healing (1–3). The fundamental dynamics of individual cell migration are generally well understood: The front of the cell extends and protrudes, generally mediated by actin polymerization. The cell adheres to the underlying extracellular matrix substrate via integrin binding and the formation of focal adhesions (4). Finally, a cell-generated contractile force retracts the cell back, generating net movement in the direction of migration (5, 6).

At the level of an individual cell, while there are many signaling pathways that regulate the dynamics of lamellipodia and filopodia cellular extensions, Rho GTPases are key drivers of this process. Rho GTPases are small GTP-binding proteins that regulate essential cellular processes and functions, including cellular adhesion, shape, and proliferation. Key members of the Rho GTPase family include Rho, Rac, and Cdc42, each of which have multiple isoforms (7). The Rho GTPase proteins cycle between active GTP-bound and inactive GDP-bound forms (8, 9). While these proteins are regulators of a host of cellular processes, a primary function of Rho GTPases is the regulation of actin cytoskeleton assembly, and thus these proteins are drivers of cellular contractility (8–11). Further, there is strong experimental evidence that Rho GTPase activity is critical for cell migration (12, 13).

As Rho GTPase activity drives actomyosin-mediated mechanical forces, mechanical forces also in turn modulate Rho GTPase activity via mechanotransduction: For example, mechanical compression of the cell membrane in human mesenchymal cells has been shown to reduce RhoA activity, mediated by mechanosensitive ion channels (14). Katsumi and colleagues show that stretch of vascular smooth muscle cells inhibited Rac and decreased lamellipodia formation (15). Thus, there is a complex and bidirectional interplay between Rho GTPase activity and mechanical forces at the cellular level (16, 17).

The *collective* migration of cells is inherently a more complex process, involving the coordination of mechanical and biochemical interactions between both cells and the surrounding substrate and neighboring cells within the tissue (10). Collective migration can be broadly described by different categories, including sheet migration, sprouting and branching, and tumor invasion (1). Coordinated migration generally requires cells to be in physical contact and coupled mechanically via cell-cell junctions. Cell-cell junction forces in turn can regulate key physiological processes, such as proliferation and differentiation (18). However, the interactions between junctional forces and collective migration dynamics are not fully understood.

In this study, we extend a prior model to develop a minimal modeling framework to investigate the interactions between mechanochemical signaling and junctional forces in collective cell migration, investigating directed migration in one dimension. We found in individual cells that migration velocity depends on the feedback between mechanical tension and Rho GTPase activity in a biphasic manner. In mechanically coupled tissue during collective cell migration, waves of Rho GTPase activity mediated mechanical contraction/extension and thus synchronization throughout the tissue. Additionally, we found that junctional forces exhibited distinct spatial patterns during collective cell migration, with larger forces near the leading edge, and further, that larger junctional force magnitudes were associated with faster collective cell migration. Our model predicts that collective cell migration depends on both the emergent dynamics and interactions between cellular-level Rho GTPase activity and contractility, and multicellular-level junctional forces.

## METHODS

In this work, we first extend a recently developed minimal cellular model coupling mechanical tension and Rho GTPase signaling to account for the front-back cellular polarity of migrating cells. We then further expand this model to account for mechanical junctions between cells and investigate the properties of collective cell migration.

### Individual cell model

The minimal mechanochemical model that serves as the framework for our study was recently presented by Zmurchok, Bhaskar, and Edelstein-Keshet (19). In this model, cellular Rho GTPase activity is coupled with cellular mechanics (Fig. 1A). Individual cell mechanics are governed by a spring-dashpot system, for which a single spring governs extension/contraction of an individual cell and two dashpots at the cell front and back represent viscous coupling with a fixed substrate (Fig. 1B). As demonstrated by Zmurchok and colleagues, in this model, mechanochemical coupling via several feedback mechanisms can drive periodic cellular contraction and extension: (1) An increase in active Rho GTPase levels promotes actomyosin-mediated cellular contraction, which in turn reduces cellular tension. (2) While Rho GTPase activity is self-activated via positive feedback, active Rho GTPase levels are subsequently reduced by the loss of cellular tension. (3) Reduced Rho GTPase activity drives cellular relaxation and extension, increasing cellular tension. (4) Increased cellular tension promotes Rho GTPase activation and increases active Rho GTPase levels, completing the cycle.

**Figure 1:**
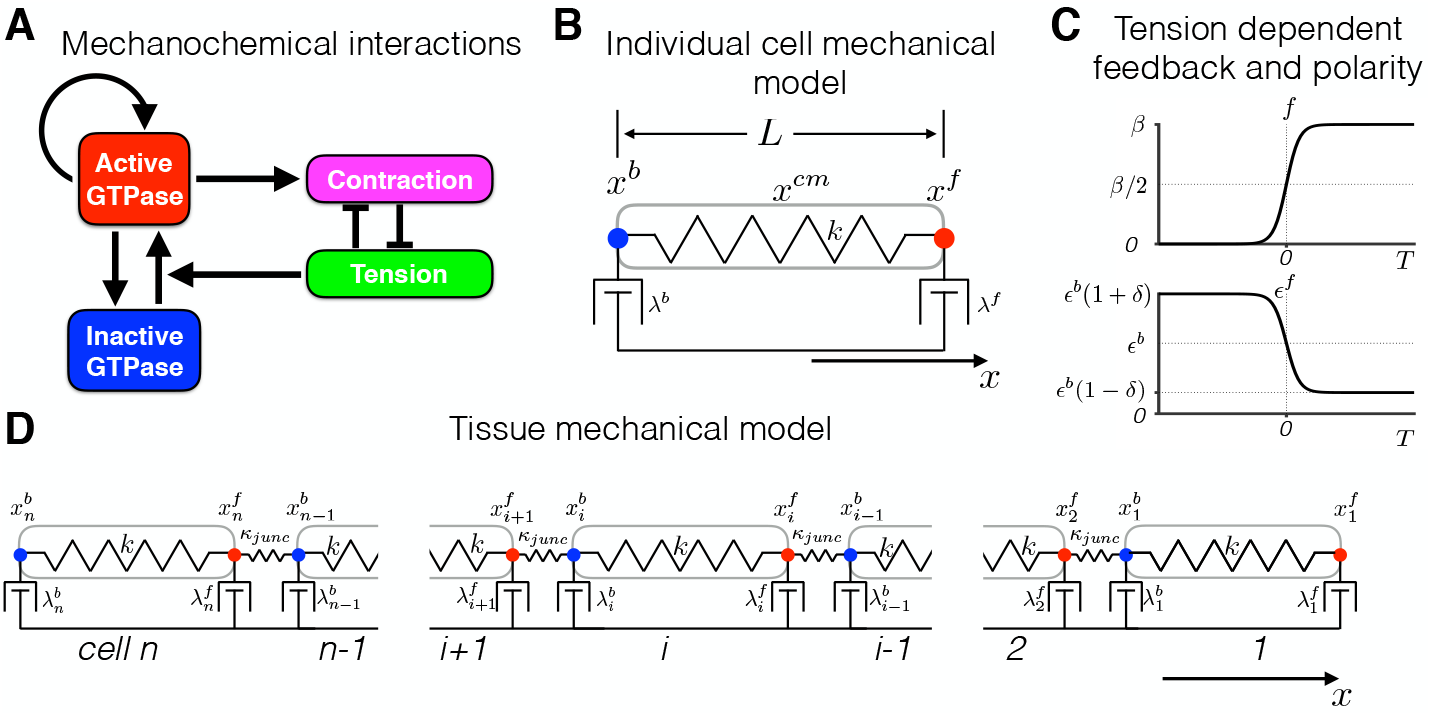
Individual cell and tissue mechanochemical models. (A) Diagram of mechaochemical interactions between active and inactive Rho GTPase levels and mechanical contraction and tension. Adapted from (19). (B) Mechanical model of an individual cell. The model governs the position of the cell front (*x*^*f*^), back (*x*^*b*^), and center of mass (*x*^*cm*^) (Eq. 1d-1e). The cell is represented by a Hookean spring, with spring constant *k*, and dashpots at the cell front and back with constants *λ*^*f*^ and *λ*^*b*^, respectively. (C, top) Tension-dependent GTPase activation *f* (*T*; *β*) (Eq. 1b) and (bottom) tension-dependent front-back polarity *ϵ*^*f*^ (*T*; *δ*) (Eq. 1f) are shown as functions of cellular tension *T*. (D) Mechanical model of *n* mechanically coupled cells. The model governs the positions of cell fronts 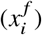 and backs 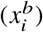 for *i* = 1, …, *n* (Eq. 2). Cell-cell mechanical junctions are represented by Hookean springs, with spring constant *κ*_*junc*_.

Normalized active Rho GTPase signaling (*G*) is governed by the following equation:

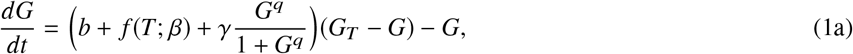

where the first term represents a tension-dependent activation rate of inactive Rho GTPase (*G*_*inactive*_ = *G*_*T*_ - *G*), *G*_*T*_ is total Rho GTPase levels (i.e., active and inactive forms), *b* is a basal activation rate, *f* (*T*; *β*) is tension-dependent activation rate, and *γ* is the Hill equation positive feedback activation rate. The second term represents a constant rate of inactivation. As in the Zmurchok model, time has been normalized relative to the active Rho GTPase residence time (or the inverse of the inactivation rate). Thus, time is presented in normalized units.

The tension-dependent activation is given by sigmoidal function of tension:

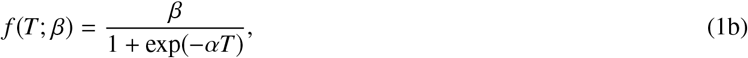

where *β* governs the strength of tension-feedback (Fig. 1C, top). Cellular tension *T* is given by the difference of cell length *L* and a Rho GTPase-dependent resting length *L*_0_, that is, *T* = *L* - *L*_0_ *G*. The function *f* (*T*; *β*) produces a switch-like response in Rho GTPase activity in the presence of cellular extension, but minimal activation for contraction.

Rho GTPase activity drives cellular contraction by reducing the cell resting length, with the following dependence:

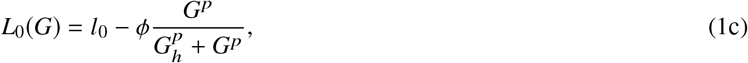

where *l*_0_ is a baseline resting length, and *ϕ* and *G*_*h*_ are Hill equation amplitude and half-maximum parameters for Rho GTPase-dependence on cell resting length, respectively. The maximum cell resting length *l*_0_ = 1; thus lengths in the spatial dimension are presented in units normalized such that the unit measure is equal to the length of a maximal relaxed cell.

Cellular mechanics are determined by a force balance at each end of the cell, with the position of the cell front *x* ^*f*^ and back *x*^*b*^ governed by the following equations, respectively:

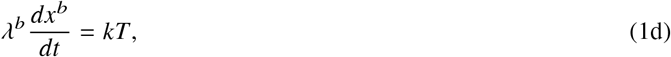

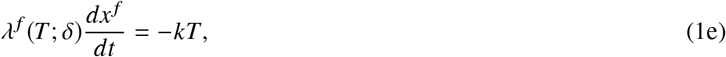

where *k* is a Hookian spring constant for the cell, and *λ*^*f*^ and *λ*^*b*^ are cell front and back viscosities or friction constants, respectively, such that cell length *L* = *x* ^*f*^ - *x*^*b*^. The cell center of mass is given by *x*^*cm*^ = (*x* ^*f*^ + *x*^*b*^)/2.

In the original model formulation by Zmurchok and colleagues, *λ* ^*f*^ = *λ*^*b*^. Here, we introduce a front-back polarity, in which the back viscosity is assumed to be fixed, given by *λ*^*b*^ = *k*/*ϵ*^*b*^, where *ϵ*^*b*^ is a normalized extension/contraction rate. However, the viscosity of the cell front *λ*^*f*^ is assumed to be tension-dependent, a formulation modified from an approach recently presented by Lopez, Das, and Schwarz (20), with *λ*^*f*^ (*T*; *δ*) = *k*/*ϵ* ^*f*^ (*T*; *δ*), where

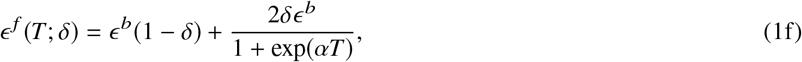

and *δ* ∈ [0, 1) is a measure of the asymmetry between the front and back of the cell (Fig. 1C, bottom). Parameter *δ* can be considered a measure of cellular front-back polarity or the strength of a guidance cue (e.g., a chemotactic gradient), where a larger *δ* corresponds to a larger polarity or gradient. The formulation is based on the assumption that during contraction while tension is low, focal adhesions at the cell front are being assembled, and thus there are fewer cell surface-integrin bonds formed, such that friction at the cell front is initially weak (resulting in a large *ϵ*^*f*^ and small *λ*^*f*^); however following focal adhesion formation, friction at the cell front is larger (i.e., a small *ϵ*^*f*^ and large *λ* ^*f*^) during the cell extension and high tension.

Note for that *δ* = 0, *ϵ*^*f*^ = *ϵ*^*b*^ and thus *λ*^*f*^ = *λ*^*b*^, as in the original model, resulting in a stationary cell. As shown in Zmurchok et al, reproduced in Fig. 2, the long-term behavior of the mechanochemical dynamics depends on the relative strength of the tension-dependent feedback: For small feedback (*β* = 0.05, Fig. 2A), the cell remains fixed in a relaxed state, while for large feedback (*β* = 0.3), the cell is fixed in a contractile state. For an intermediate feedback (*β* = 0.16), the cell state oscillates between contraction and extension, coupled with oscillations in Rho GTPase levels. Note that while the position of the cell front *x*^*f*^ and back *x*^*b*^ extend and contract, the center of the cell *x*^*cm*^ remains constant, as expected for *δ* = 0. Fig. 2B demonstrates the transition from relaxed (red), to oscillatory (black), to contractile (blue) states, as a function of *β*. Further, we find that in the oscillatory regime, the period of oscillations also depends on the strength of the tension-dependent feedback, such that the period increases as *β* increases (Fig. 2C)

**Figure 2:**
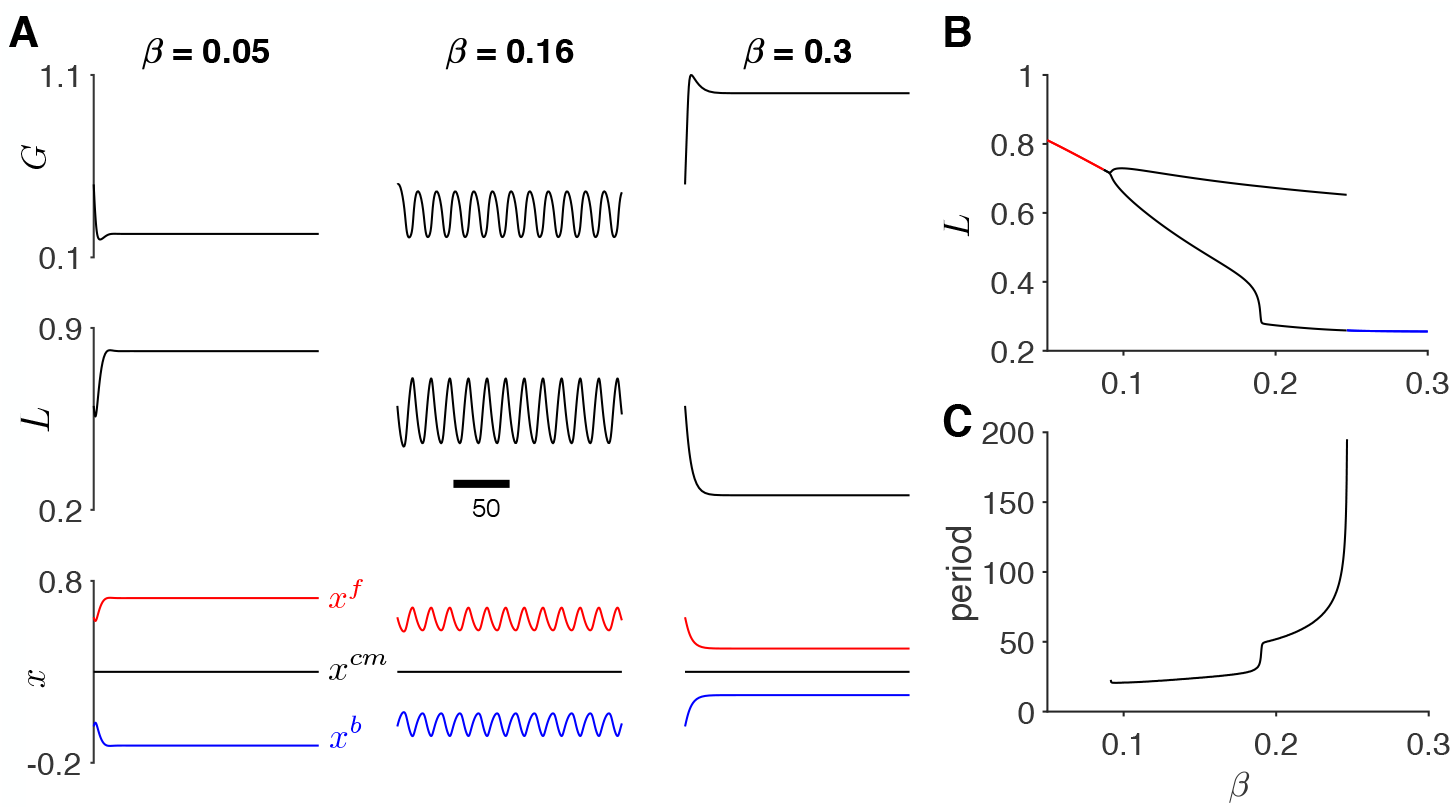
Dynamics of a stationary individual cell. (A) The time course for Rho GTPase activity *G*, cell length *L*, and cell position (front (red) *x*^*f*^, center (black) *x*^*cm*^, and back (blue) *x*^*b*^) are shown for low, intermediate, and high feedback-tension parameter *β*. For low *β*, cellular Rho GTPase activity is low and the cell is a fixed relaxed state. For high *β*, Rho GTPase acitivity is high, and the cell is in a fixed contractile state. For intermediate *β*, Rho GPTase activity oscillates, with corresponding periodic cellular extension and contraction. (B) The steady-state length is shown as a function of *β*, demonstrating the relaxed (red), oscillatory (black), and contractile (blue) regimes. (C) In the oscillatory regime, oscillation period increases as a function of *β*, with steep dependence near the contractile regime. Parameters: *δ* = 0.

### Collective cell migration model

We next extend the minimal migrating cell model to represent a 1-dimensional tissue of mechanically coupled cells (Fig. 1D). For each cell *i*, Rho GTPase activity (*G*_*i*_) and mechanical forces are represented; in addition to cellular tension, an intercellular mechanical junction between the back of cell *i* and the front of cell *i* + 1 is represented by a Hookian spring with spring constant *κ*_*junc*_ and resting length *L*_*junc*_. Thus, the tissue model of *n* cells is represented by Rho GTPase activity (*G*_*i*_), and the position of the cell front 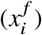 and back 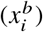, i.e., 3*n* variables, governed by

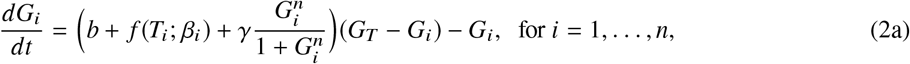

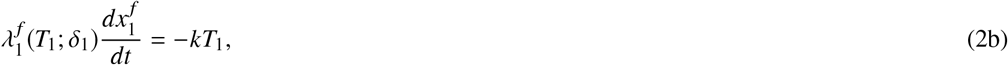

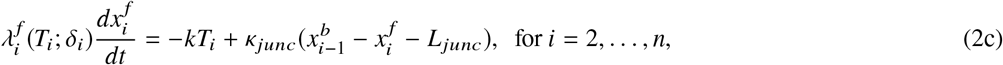

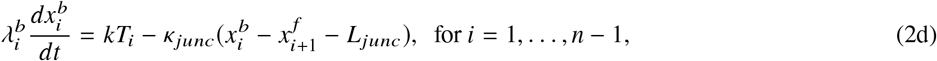

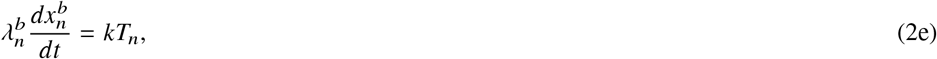

where cell length 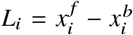, and cell tension *T*_*i*_ = *L*_*i*_ - *L*_0_(*G*_*i*_). We assume that cells may have different parameters for front-back polarity (*δ*_*i*_) and tension feedback (*β*_*i*_). We note that Zmurchok and colleagues also consider a 1-dimensional array of cells, but they did not represent intercellular mechanical forces transmitted via a spring, but rather assumed that the back of cell *i* and the front of cell *i* + 1 are the same “node,” which would be the limit of *L*_*junc*_ = 0 and *κ*_*junc*_ → ∞. Importantly, our formulation enables explicit prediction of intercellular junctional forces throughout the tissue.

All model parameters are given in Table S1. In all individual cell or tissue simulations, initial conditions for Rho GTPase activity *G* 0 = 1. Initial cell positions were defined such that initial cell lengths were equal to 0.6, with intercellular node spacing such that mechanical junctions were in equilibrium (i.e., front-back cell position differences were equal to the junctional resting length *L*_*junc*_). Unless otherwise noted, simulations were run for a duration of 10,000 normalized time units, and analysis was performed on the last 5,000 time units.

## RESULTS

### Individual cell migration depends on tension-dependent feedback and front-back polarity

We first demonstrate how the introduction of front-back cell polarity produces cell migration in an individual cell (Fig. 3A). For *β* in the oscillatory range and a small front-back polarity (*δ* = 0.1), cell front (red) and back (blue) demonstrate periodic extension and contraction (as in the stationary cell case shown in Fig. 2). However, the asymmetry in the cell front friction (specifically more friction, i.e., *λ*^*f*^ > *λ*^*b*^, during contraction and less friction, i.e., *λ*^*f*^ < *λ*^*b*^, during extension) produces a gradual net movement in the positive *x* direction. The position of the cell center *x*^*cm*^ (black) illustrates the gradual migration as well. The time course of the cell front, center, and back illustrate that as *δ* increases, cell migration velocity increases (see Movies S1 and S2 in Supporting Material).

**Figure 3:**
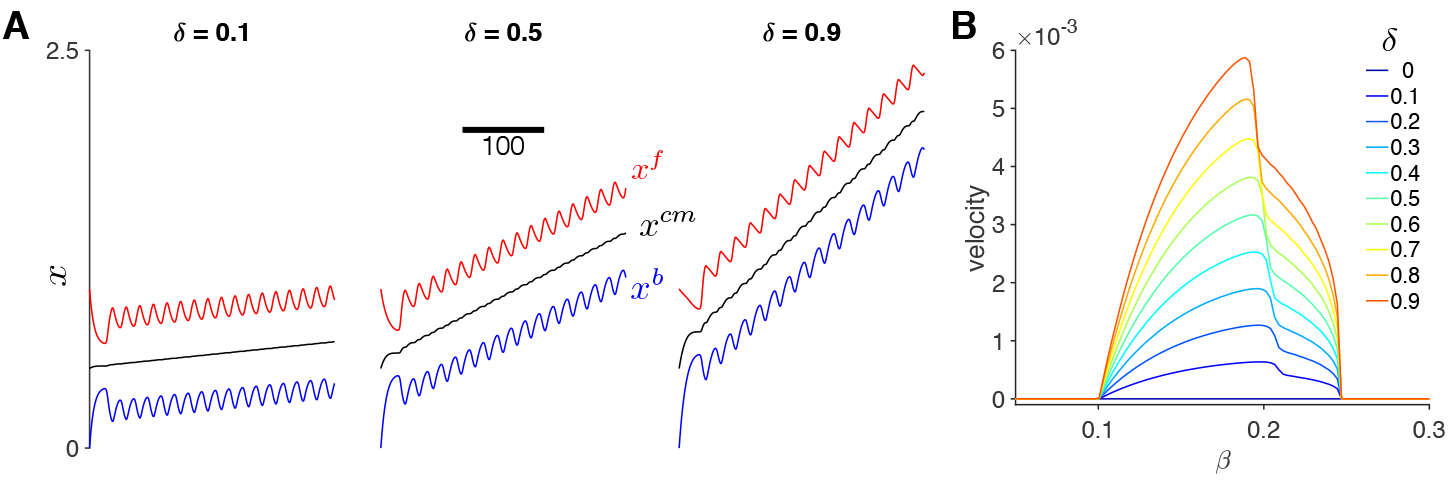
Dynamics of a migrating individual cell. (A) The time course for cell position (front (red) *x*^*f*^, center (black) *x*^*cm*^, and back (blue) *x*^*b*^) are shown for different values of front-back polarity parameter *δ*. Cell migration velocity increases for increasing *δ*. (B) Cell migration velocity is shown as a function of tension-feedback parameter *β* for different values of *δ*. Velocity is zero in the non-oscillatory *β* regime, and a biphasic function of *β* in the oscillatory *β* regime. Parameters: (A) *β* = 0.16

In Fig. 3B, we plot the cell migration velocity as a function of *β* for different values of *δ*. Velocity is calculated as the slope from a linear regression on the cell center position *x*^*cm*^ versus time (using the second half of the simulation for analysis). We find that for any value of *β*, increasing *δ* increases migration velocity (consistent with Fig. 3A). Additionally, the model predicts that cell migration requires periodic extension and contraction, and that cells fixed in a relaxed or contractile state are stationary. Thus, the *β* parameter regimes for oscillations are identical to the regime for non-zero cell migration velocities, and for *β* values outside the oscillation regime, migration velocity is zero. We also find that for a given value of *δ*, velocity is biphasic as a function of *β*, such that there is an optimal *β* that yields the fastest migration velocity. The optimal *β* value moderately depends on *δ*, decreasing as *δ* increases, demonstrating feedback between cell tension and front-back polarity; however, the *β* value for the fastest single cell migration velocity is between 0.19 and 0.2 for the full range of *δ* values.

### Collective cell migration depends on individual cell properties and junctional forces

After determining the relationship between tension-dependent feedback and front-back polarity on migration in an individual cell, we next consider these interactions on collective cell migration on mechanically coupled cells. We investigate both homogeneous tissues, in which all cells have the same individual properties, and heterogeneous tissues, in which the front or “leader” cell may have different properties than the back or “trailing” cells in the tissue.

### Homogeneous tissues

We first consider the case of a homogeneous tissue comprised of 10 cells, in which *β* is in the oscillatory regime for all cells, but no front-back polarity (*δ* = 0, Fig. 4). As expected, the collective tissue is stationary; however, individual cells in the front and back of the tissue exhibit periodicity around a stationary position (Fig. 4A). Thus, cells in the front (e.g., cell 1) and back (e.g., cell 10) exhibit both the higher frequency oscillations, corresponding to periodic extension and contraction, and a slower frequency oscillation that is an emergent property of the tissue. Towards the interior of the tissue (e.g., cells 4 and 5), oscillations due to extension/contraction are substantially dampened. In this stationary tissue, there is also a symmetry in the time course of the cell positions, in that the front of cell 1 mirrors the back of cell 10, the front of cell 2 mirrors the back of cell 9, etc, as might be expected in this symmetrical tissue.

**Figure 4:**
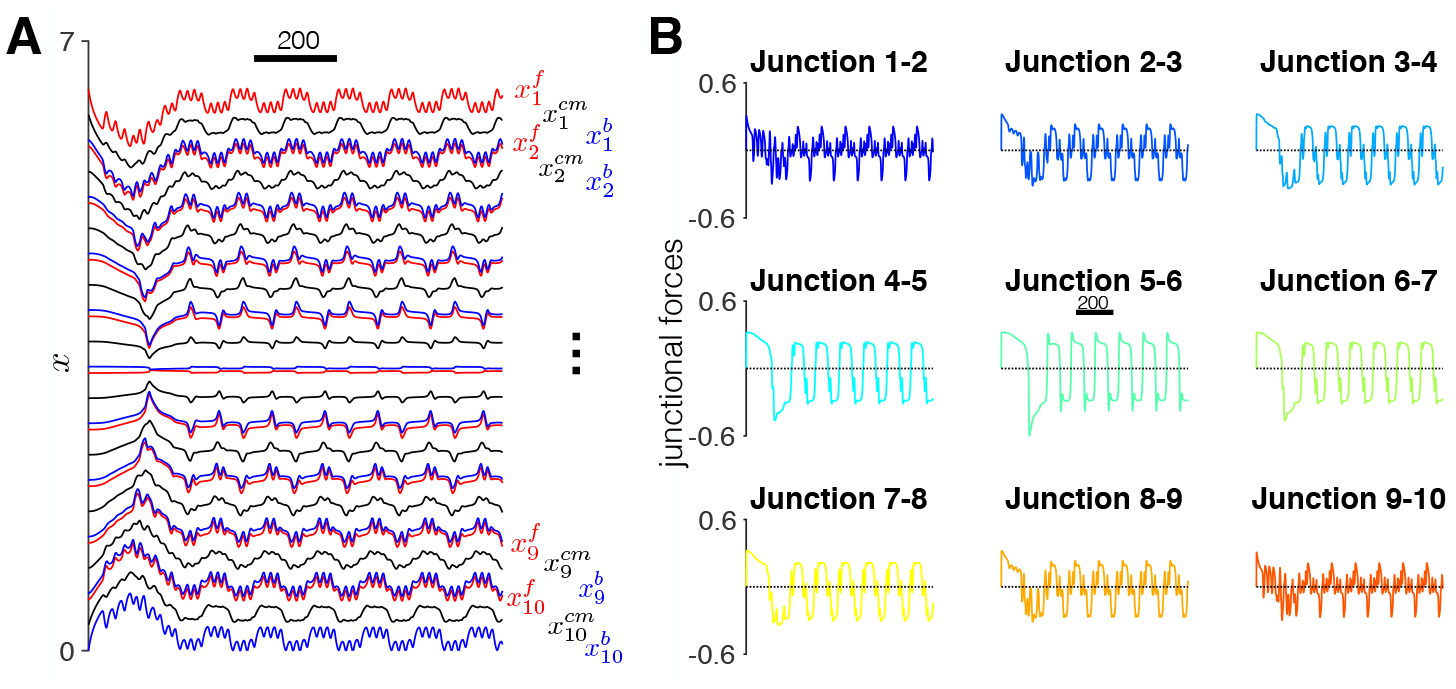
Dynamics of a stationary tissue. (A) The time course for cell positions (front (red) 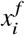, center (black) 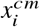, and back (blue) 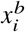, for *i* = 1, 2, …, *n* = 10) are shown. The time courses exhibit a front-back symmetry and two distinct frequencies, corresponding to the individual cellular contraction and extension and an emergent frequency from cell-cell mechanical coupling. (B) The time course for each cell-cell junction also exhibit multiple frequencies and periods of both tension and compression. Parameters: *β* = 0.16, *δ* = 0.

In Fig. 4B, we investigate the time course of the junctional forces between each cell pair (e.g., between cell 1 and 2, between cell 2 and 3, etc). The time course for junctional forces demonstrates that the cell junctions exhibit time periods of both tension and compression. Junctional forces on the periphery of the tissue (e.g., junction 1-2 and junction 9-10) also exhibit periodicity associated with both contraction and extension and a slower component associated with tissue movements, as in the cell position plots. The higher frequency associated with cell contraction and extension is similarly dampened in the tissue interior (e.g., junction 5-6). Thus, the stationary tissue simulations predict two key properties: emergent tissue-scale dynamics and a spatial pattern for junctional forces. To investigate the mechanism of the emergent frequency in tissue, we plot Rho GTPase activity in each cell in Fig. 5 and S1. We find that for cells on the periphery (e.g., cell 1 and 10), Rho GTPase activity oscillations are entrained with the cellular contractions and extensions. However, Rho GTPase activity in the interior (e.g., cell 5 and 6) is entrained to the lower frequency oscillations. Further, we observe Rho GTPase activity waves that propagate from the tissue periphery to the interior (dashed arrows). Thus, our simulations predict that the tissue-scale emergent dynamics arise due to the interactions between cell-cell coupling and cellular Rho GTPase activity (see Movie S3 in Supporting Material).

**Figure 5:**
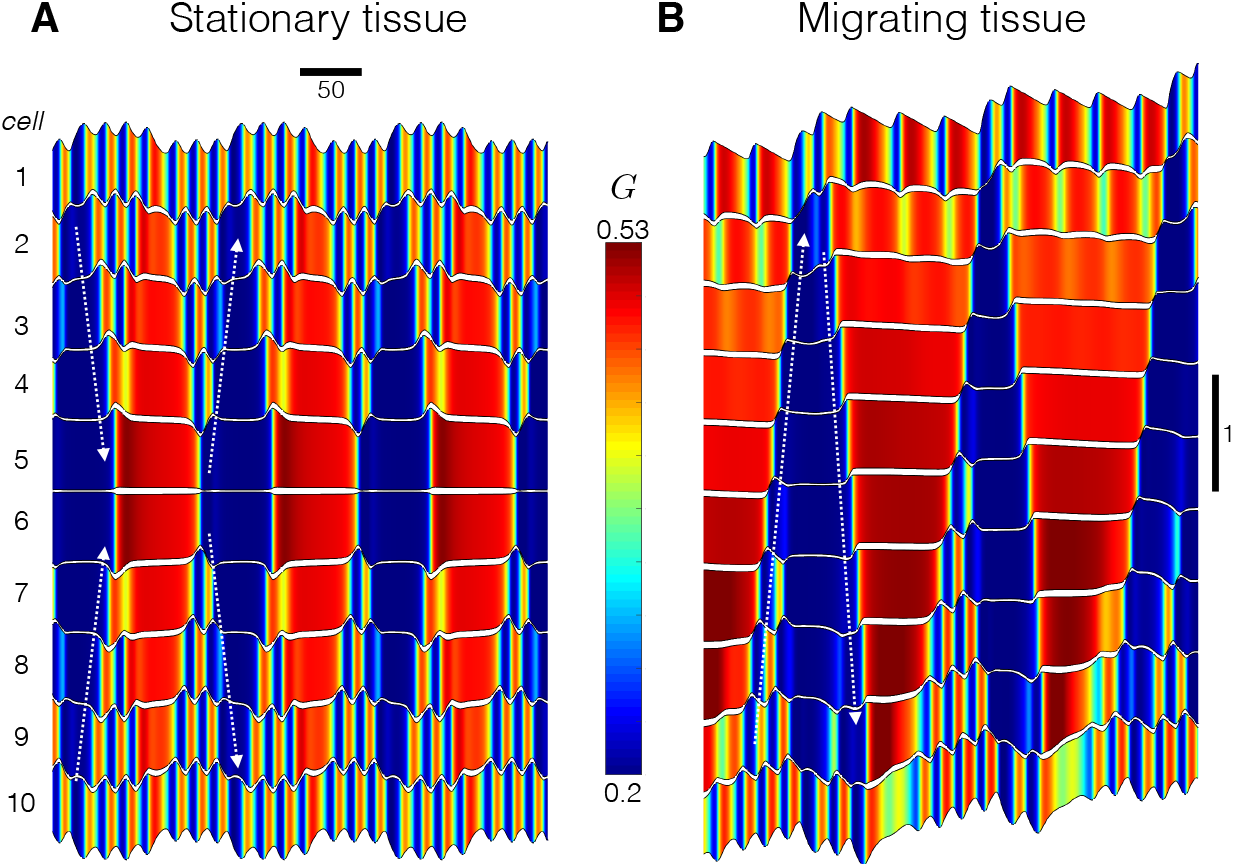
Dynamics of Rho GTPase activity in homogeneous tissue. Kymographs show the time course for cell positions (black), and color indicates Rho GTPase activity in each cell (*G*_*i*_, for *i* = 1, 2, …, *n* = 10) in (A) stationary and (B) migrating tissue. Parameters: *β* = 0.16, (A) *δ* = 0, (B) *δ* = 0.9.

We next investigate simulations of homogeneous tissue in which individuals cells exhibit a front-back polarity (Fig. 6). As in the stationary tissue, the time course of cell positions exhibits complex emergent dynamics: specifically multiple frequencies, a faster frequency associated with cell contraction and extension and a slower frequency from tissue movement that emerges due to cell-cell coupling (Fig. 6A). Further, as in individual cells, increasing the front-back polarity (i.e., increased *δ*) increases the velocity of the collective cell migration. In Fig. 5B and S1B, we plot Rho GTPase activity in the migrating tissue. As in the stationary tissue, Rho GTPase activity in the periphery exhibits high frequency oscillations associated with cell contraction and extension, while the interior exhibits slower frequency oscillations that are an emergent property of the tissue. Further, we observe Rho GTPase activity waves propagating in the tissue. However, in contrast with the stationary tissue, we observe low Rho GTPase activity wave propagate from tissue back to front, and then high Rho GTPase waves propagating front to back (dashed arrows, see Movie S4 in Supporting Material). The timing of these Rho GTPase activity waves correspond to the beginning and end of the emergent slower frequency, facilitating synchronization across the tissue, again demonstrating that interactions between cell-cell coupling and Rho GTPase activity drive emergent tissue properties.

**Figure 6:**
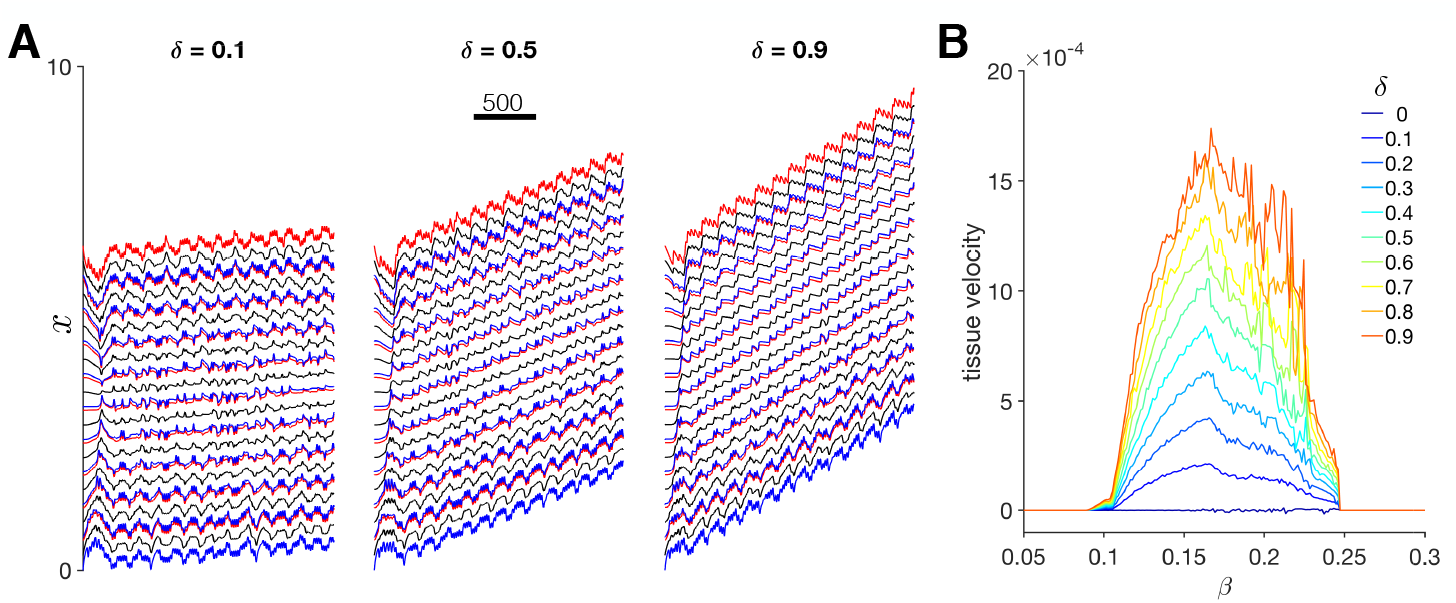
Dynamics of collective cell migration in homogeneous tissue. (A) The time course for cell positions (front (red) 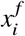, center (black) 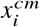, and back (blue) 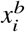, for *i* = 1, 2, …, *n* = 10) are shown for different values of front-back polarity parameter *δ*. Collective cell migration velocity increases for increasing *δ*. (B) Tissue migration velocity is shown as a function of feedback-tension parameter *β* for different values of *δ*. Velocity is zero in the non-oscillatory *β* regime, and a generally biphasic but jagged function of *β* in the oscillatory *β* regime. Parameters: (A) *β* = 0.16.

We summarize the tissue velocity as a function of *β* for a range of *δ* values (Fig. 6B). Velocity of the tissue is calculated as the slope from the linear regression of the first cell center 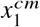 versus time. We find several key similarities and differences between individual cells and tissues: As in individual cells, collective cell migration occurs only for *β* values within the oscillatory regime, and also similarly, velocity increases as *δ* increases. Further, we find that for a given *β* and *δ* parameter combination, migration velocity of the homogeneous tissue is slower compared to the corresponding individual cell. The general shape of the velocity dependence on *β* is also biphasic, as in individual cells. However, the velocity versus *β* traces are “noisy” or jagged, in comparison with the smooth traces for individual cells (cf. Fig. 3B), and this jaggedness is more pronounced for larger *δ*. This suggests that small changes in cell contractility (i.e., changes in *β*) can result in quite large changes in collective cell migration velocity, which emerges due to forces imposed at the cell-cell junctions that desynchronize oscillations of mechanically coupled cells, discussed in more detail later. In Supporting Material, we plot the tissue velocity against the corresponding individual cell velocity, for a given value of *β* and *δ* (Fig. S2) and show that, while in general the tissue and individual cell velocities are proportional, that is not the case for all values of *β*.

We next investigate junctional forces in collective cell migration (Fig. 7). As in the stationary tissue, the time course of junctional forces exhibits periods of both tension and compression; however, in contrast, forces are not symmetric (Fig. 7A). We compute the average junctional force for each cell-cell junction and find that junctional forces at the front of migrating tissue are on average tensile, while junctional forces at the back of the tissue are on average compressive (Fig. 7B). The junctional force average exhibits a distinct spatial pattern, with largest forces near, but not at, the tissue front, and a gradual decrease towards the tissue back.

**Figure 7:**
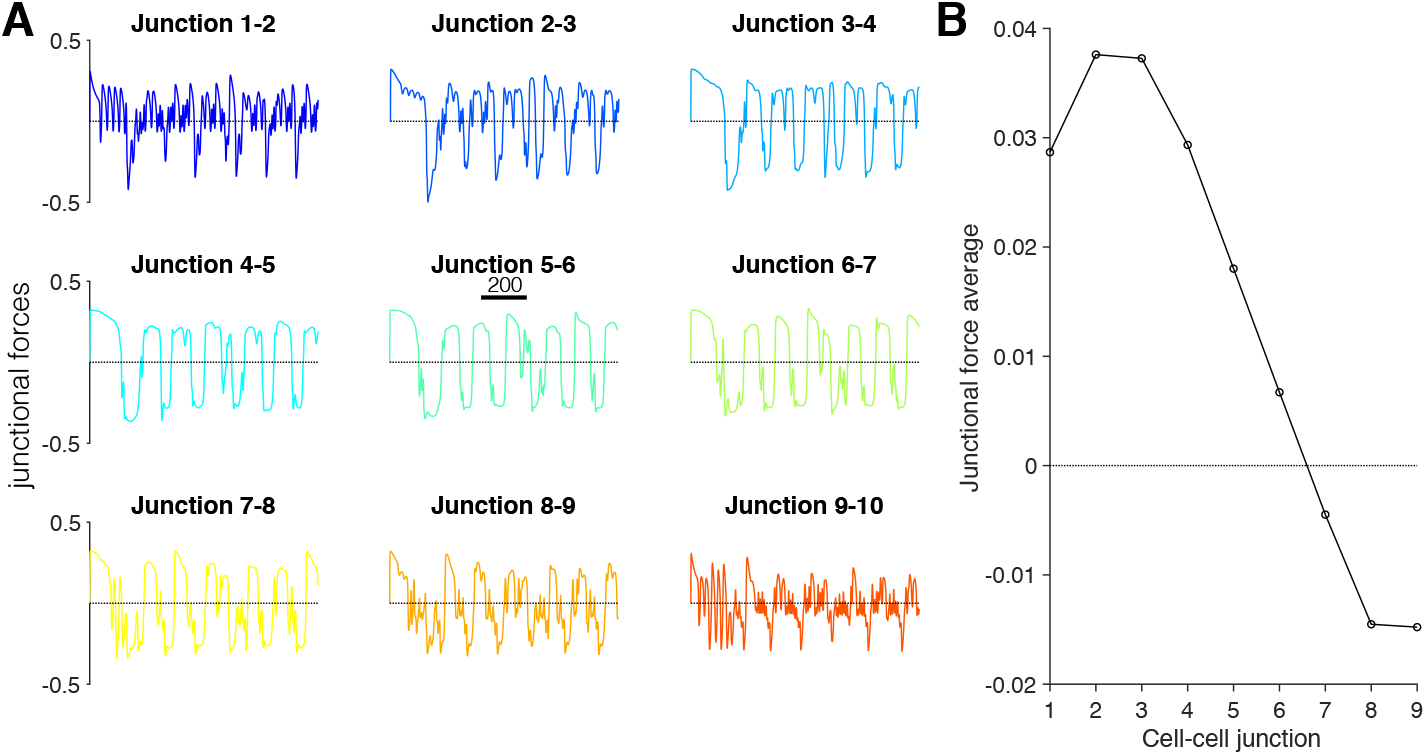
Spatial pattern of junctional forces in collective cell migration. (A) The time course of each cell-cell junction exhibits multiple frequencies, in particular at the tissue periphery. (B) The junction force average is shown as a function of the cell-cell junction (with 1 corresponding to the cell 1-cell 2 junction at the tissue front). The junctional force average exhibit a maximum near, but not at, the tissue front, and then decreases towards the tissue back. Parameters: *β* = 0.16, *δ* = 0.5

In Fig. 8A, we plot cell-cell junctional force averages as a function of *β* for different values of *δ*. We find that for all migrating tissues (i.e., *δ* > 0), we find the same general spatial pattern for all values of *β*: largest force near, but not at the tissue front, and decreasing towards the tissue back. Further, the magnitude of junctional force averages are generally larger for tissues with faster velocities (cf. Fig. 6B): the junctional forces are generally “jagged” biphasic functions of *β* and junctional force magnitudes increase for increasing *δ*. A scatter plot of the junctional force averages for the different cell-cell junctions plotted against the corresponding tissue velocity for all values of *β* and *δ* further demonstrate that larger magnitude junctional forces in the tissue front and back are associated with faster tissue velocities (Fig. 8B).

**Figure 8:**
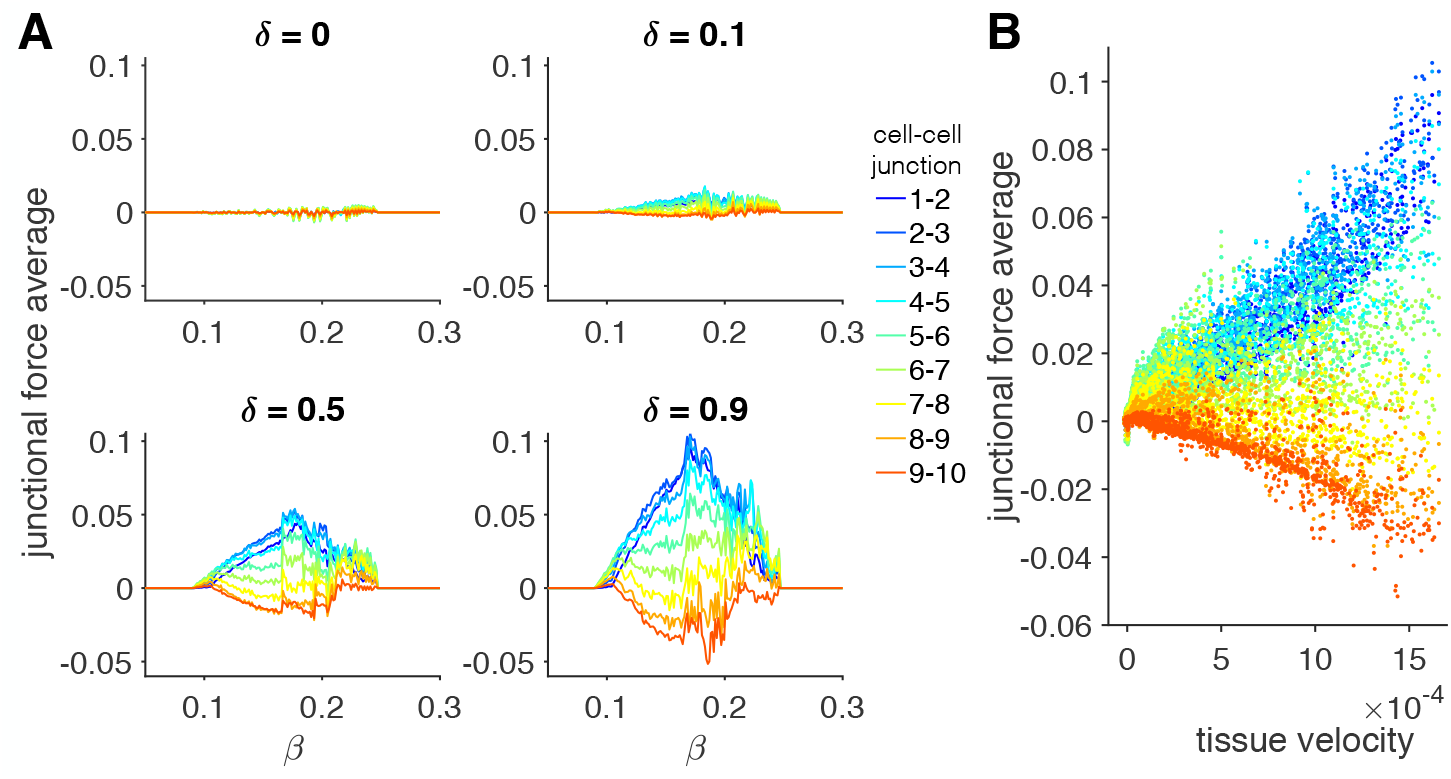
Spatial pattern of junctional forces depends on tension-feedback and front-back polarity. (A) Junctional force averages are shown for all cell-cell junction pairs as a function of *β* for different values of *δ*. For migratory tissues (*δ* > 0), junctional force averages decrease in general decrease from the tissue front (cell 1-cell 2 junction) to the tissue back. The magnitude of junctional force averages generally increases with increasing *δ* and follows a jagged biphasic dependence on *β*. (B) A scatter plot of the junctional force averages for different cell-cell junction pairs (different colors) against the corresponding tissue velocity, calculated from simulations for varying *β* and *δ* demonstrate that larger magnitude junctional forces occur for faster collective cell migration.

We next investigate the mechanism of steep sensitivity to the feedback-tension (*β*) in collective cell migration. In Fig. 9, we consider two tissues, with *β* values differing by approximately 1%. For *β* = 0.22588 (Fig. 9A), we find that the cell positions time course is comparable to the examples in Fig. 6A, with oscillations corresponding to cell contraction and extension and slower emergent frequent from mechanical coupling between cells. Further, the time courses also reveal mechanical “waves” that propagate from the tissue back to front and from front to back that arise due to synchronization of individual cell extensions (denoted by black arrows). However, there also several instances in which the mechanical waves “fail” and do not propagate to the tissue front (denoted by orange block symbol), due to desynchronization between the individual cells.

**Figure 9:**
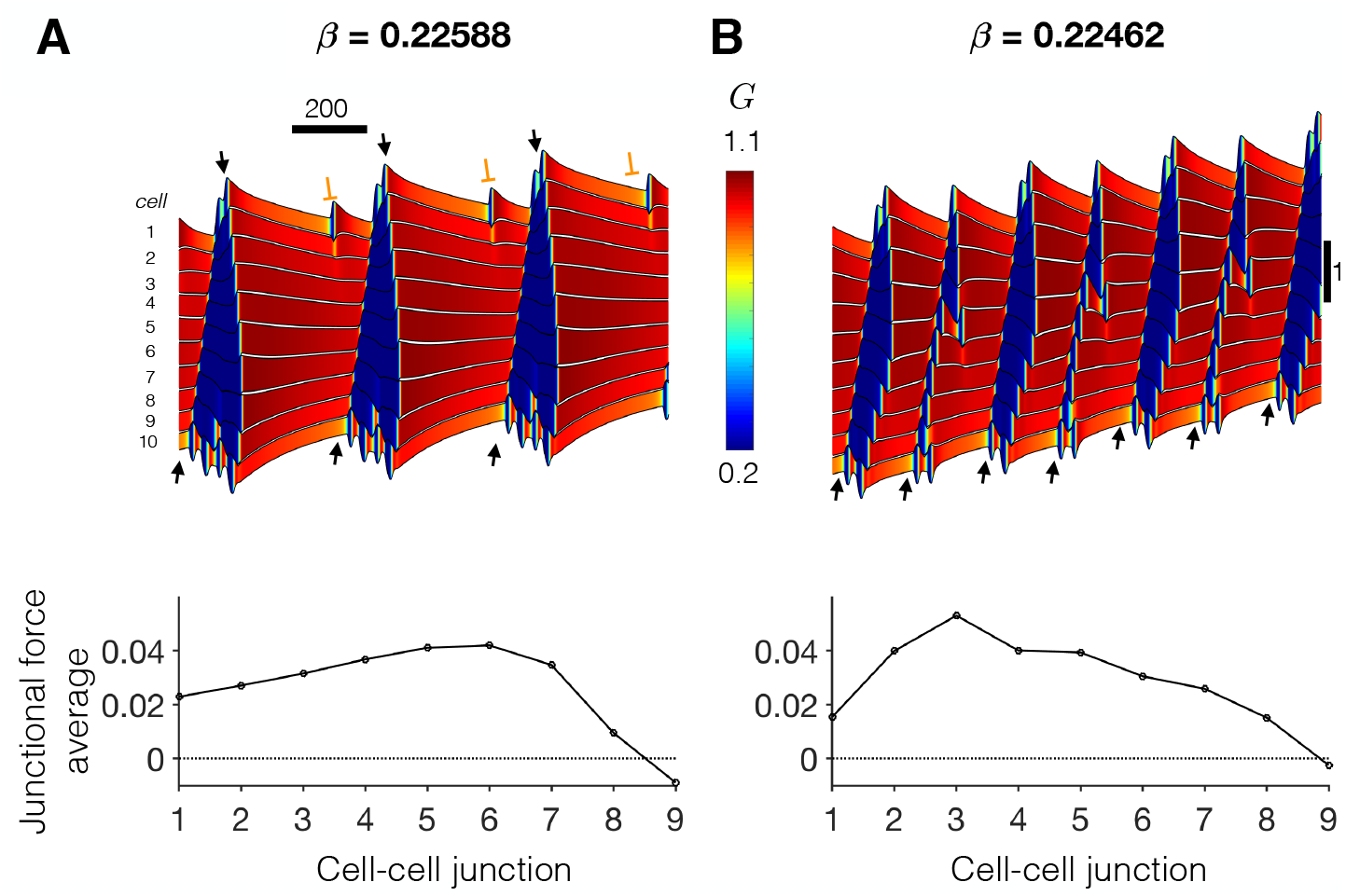
Mechanochemical resonance during collective cell migration. Kymographs show the time course for cell positions (black), and color indicates Rho GTPase activity for two values of tension-feedback parameter *β* (A, B) that differ by approximately 1%. Successful and failed mechanical wave propagation are denoted by black arrows and orange block symbols, respectively. (Bottom) The spatial pattern of junctional force averages for the two examples. Parameters: *δ* = 0.9.

For a slightly smaller *β* = 0.22462 (Fig. 9B), the oscillation period of individual cells is shorter (note the steep dependence of the period on *β* in Fig. 2C), such that after an initial transient, all mechanical waves successfully propagate from back to front. Rho GTPase activity facilitates a tissue-wide synchronization such that the emergent tissue frequency is nearly twice that for the slightly larger *β* value. This synchronization can be described as a “mechanochemical resonance,” in which the individual cell oscillation frequency more closely matches the emergent tissue frequency that results in an increased collective cell migration velocity (see Movie S5 in Supporting Material). As such, the 1% change in *β* results in an approximately 51% change in velocity (0.001142 vs. 0.0005770). The junctional force average spatial patterns are similar to previous examples; however for the the slower migrating tissue, the largest junctional forces are more towards the tissue interior.

Summary plots of tissue velocity for tissues of different sizes are shown in Supporting Material (Fig. S3A). Tissue velocity in general decreases as the size of the tissue increases. We also find the same general trends in all tissue sizes: velocity increases as *δ* increases, and there is a jagged biphasic dependence on *β*, although the “peak” *β* value differs somewhat for different tissue sizes. We also find the same general junctional force average spatial patterns in tissues of different sizes (Fig. S3B), with the largest forces near, but not at, the tissue front. Additionally, larger magnitude forces are observed in larger tissues.

### Heterogeneous tissues

We next investigate collective cell migration in heterogeneous tissues. Specifically, we consider the case in which the properties of the cell at the front of the tissue, the lead cell, has different properties than the other 9 trailing cells towards the back of the tissue. Further, we are interested in the case in which the lead cell would individually migrate faster compared with the trailing cells individually (see Fig. 3B). For simplicity, we limit our study to cases in which either *β* or *δ* is the same throughout the tissue. If we consider a single cell simulation with (*β, δ*) parameters corresponding to a migration velocity near the peak of the biphasic curves in Fig. 3B, it should be clear that migration velocity may be slower from one of three possible parameter changes: (i) decreasing *β* (keeping *δ* constant); (ii) increasing *β* (keeping *δ* constant); or (iii) decreasing *δ* (keeping *β* constant).

We show an example of heterogeneous tissue migration in Fig. 10. Parameters for the lead cell are *β* = 0.16 and *δ* = 0.9, which an individual cell with corresponding parameters would have a velocity of approximately 0.005018. Parameters for the 9 trailing cells are such that individual cells would have a migration velocity of 20% of the lead cell individual velocity, approximately 0.001004. In Fig. 10A, the trailing cells parameters are smaller *β* = 0.1082 and the same *δ* = 0.9, and in Fig. 10B, the trailing cells parameters are a larger *β* = 0.2454 and the same *δ* = 0.9, compared with the lead cell. In both of these examples, the time series of cell positions illustrates that cell contractions and extensions are less pronounced, compared with the homogeneous tissues. In Fig. 10C, the trailing cells parameters are the same *β* = 0.16 and a smaller *δ* = 0.1885. The cell positions exhibit larger cell contractions and extensions, compared with the low and high *β* examples, but smaller compared with homogeneous tissues.

**Figure 10:**
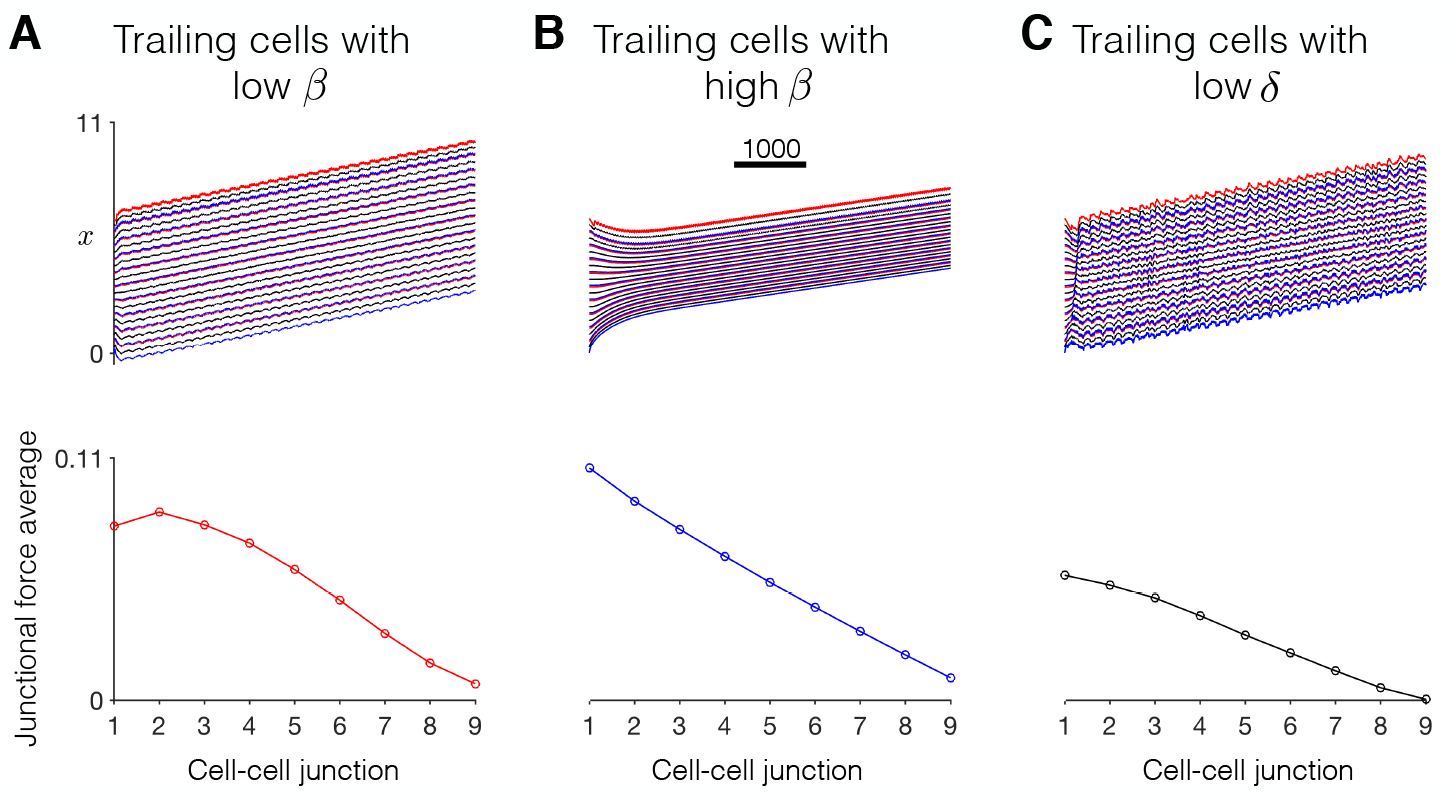
Dynamics of collective cell migration in heterogeneous tissue. (Top) The time course for cell positions (front (red) 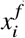, center (black) 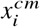, and back (blue) 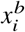, for *i* = 1, 2, …, *n* = 10) are shown for different trailing cell properties, relative for to the lead cell, with (A) low *β*, (B) high *β*, and (C) low *δ* trailing cells. See text for details. (Bottom) Spatial pattern of junctional force averages as a function of the cell-cell junction. Parameters: Lead cell: *δ* = 0.9, *β* = 0.16. Trailing cells scaling factor of 0.2: (A) *δ* = 0.9, *β* = 0.1082, (B) *δ* = 0.9, *β* = 0.2454, (C) *δ* = 0.1885, *β* = 0.16.

Interestingly, we find that the heterogeneous tissue with low *β* trailing cells has the fastest migration velocity, the tissue with low *δ* next fastest, and the tissue with high *β* the slowest. These differences are likely due to the different intrinsic oscillation periods and amplitudes of the trailing cells in each case. For the trailing cells with low *β*, the amplitude and period of the individual trailing cells oscillations are relatively close to the lead cell such that a larger degree of mechanical synchronization occurs and a faster resulting tissue velocity. In contrast, for high *β* trailing cells, the amplitude and period of the trailing cells oscillations differ dramatically from the lead cell, resulting in mechanical desynchronization and a slow tissue velocity. In the tissue with low *δ* trailing cells, although the individual cells have the same intrinsic period, the different front-back polarities introduce some degree of desynchronization, similar to as observed in the homogeneous tissue. The spatial pattern of junctional force averages is similar to previous examples, with the largest forces at or near the tissue front and decreasing towards the tissue back. However, we also find that average junctional forces are positive at the tissue back, in contrast with the homogeneous tissue example in Fig. 7B. Additionally, we find a less clear correlation between junctional forces and velocity in the heterogeneous tissues, in contrast with homogeneous tissues. The tissue with high *β* trailing cells, i.e., the slowest tissue, exhibits the largest junctional force averages, while the tissue with low *δ* trailing cells, i.e., the second fastest tissue, exhibits the smallest junction force averages.

In Fig. 11, we summarize a wide range of similar heterogeneous tissue simulations, measuring tissue velocity and junctional forces as a function of the trailing cell velocity scaling factor (e.g., 0.2 in Fig. 10) for trailing cells with low *β*, high *β*, and low *δ*, relative to the lead cell. For a lead cell with *β* = 0.16 and *δ* = 0.5 (Fig. 11A), tissue velocity increases as the trailing cell velocity scaling factor increases. That is, as might be expected, as the velocity of the individual trailing cells increases and becomes closer to the lead cell velocity, collective cell migration is faster. As in Fig. 10, collective cell migration is faster for trailing cells with low *β*. Velocity is similar for low *δ* trailing cells, and velocity is slower for high *β* trailing cells. We also note that for trailing cell velocity scaling factor of 0, the trailing cells individually are stationary; however the collective cell migration velocity is still non-zero for all three cases. Thus, our model predicts that a single migratory lead cell can drive collective cell migration of a tissue comprised primarily of non-migratory cells.

**Figure 11:**
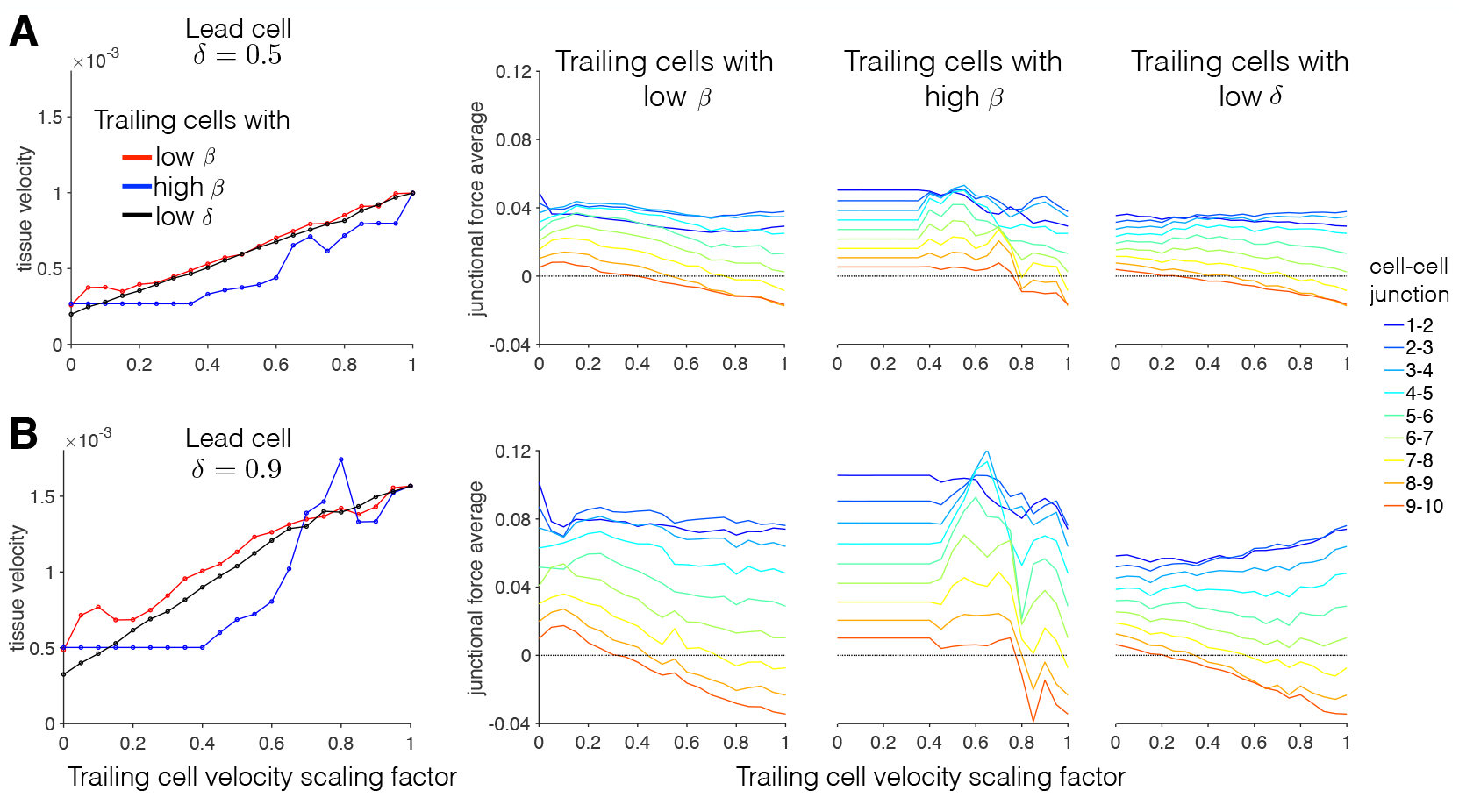
Summary of heterogeneous tissue cell migration properties. (Left) For lead cells with *δ* = (A) 0.5 and (B) 0.9, tissue velocity is shown against the trailing cell velocity scaling factor, for low *β* (red), high *β* (blue), and low *δ* (black) trailing cells. See text for details. (Right) Junction force averages are shown for all cell-cell junction pairs as a function of the trailing cell velocity scaling factor. Parameters: Lead cell: *β* = 0.16. Tissue size of 10 cells.

Spatial patterns of junctional forces for different trailing cell velocity scaling factors demonstrate the junctional force decrease from the tissue front to back, as described above. As in Fig. 10, for small scaling factors, junctional forces are positive at the tissue back. As the scaling factor increases, the magnitude of junctional forces at the tissue front remains fairly constant, while junctional forces at the tissue back decrease and become negative for larger scaling factors. Noting that this change in junctional forces occurs in conjunction with increasing tissue velocity suggests that negative average junctional forces at the tissue back promotes faster migration.

In Fig. 11B, we consider a lead cell with *δ* = 0.9. We observed similar trends as in panel A, with larger velocities and larger magnitude junctional forces, consistent with trends in the homogeneous tissue in Fig. 6 and 8, respectively. Interestingly, we find that for high *β* trailing cells, while tissue velocity is generally slow for small scaling factors, for larger scaling factors, tissue velocities are faster than the homogeneous tissue (scaling factor of 1), demonstrating that mechanical synchronization within the tissue of slower individual cells can result in faster migration than weaker synchronization of faster individual cells.

## DISCUSSION

### Summary of main findings

In this study, we present a minimal framework for modeling the dynamics of Rho GTPase-mediated migration of individual cells and tissue in one dimension. Simulations in an individual cell demonstrate that periodic Rho GTPase activity is critical for migration (Fig. 3), consistent with experiments. We find both similarities and differences between individual and collective cell migration. While mechanical coupling of *individual* cells with fast migration velocity generally corresponds to faster *collective* migration, our model predicts that tissue migration velocity is not simply governed by individual migration velocities alone (Fig. 6 and S2). We find that collective migration is governed by propagating waves of Rho GTPase activity that synchronize mechanical waves. As a result, our model predicts high sensitivity to the individual cell tension feedback (Fig. 9), in which small changes can result in large differences in tissue synchronization. We also find that collective migration exhibits a distinct spatial pattern for junctional forces (Fig. 7), with larger forces near the leading edge of the tissue, and further that larger junctional forces correspond with faster collective migration and large tissues. Finally, we predict in heterogeneous tissues that a single migratory leader cell can drive migration of a tissue comprised of slowly migrating or stationary trailing cells, in a manner that is highly dependent on the properties of the trailing cells (Fig. 10).

### Prior computational studies of cell migration

There are many prior computational studies of individual and collective cell migration, incorporating a wide range of biophysical detail (20–30). For example, Zaman and colleagues minimally model individual cell migration, integrating a force-based model of an individual cell with the interactions between cellular and extracellular mechanical forces and extracellular matrix signaling (21–23). Other studies have considered a more spatially detailed physical representation of mechanical interactions between individual cells and the extracellular matrix (25, 26). Lopez et al utilized the same one-dimensional spring representation as we consider here, with cell contraction and extension driven by hysteresis in the cell spring resting length and cell front viscosity terms (20). Vertex-based models have been developed to study collective migration in a cell sheet or tissue, modeling tissue-scale dynamics such as morphogenesis, which consider mechanical force-balance at each node along the boundary of cells and accounting for active and passive mechanical forces (31–33). Szabo and colleagues integrated individual cell mechanics into a Cellular Potts framework to predict collective migration in cell sheets (34).

While these studies focused primarily on the dynamics of cellular and extracellular mechanical interactions, other studies of cell migration have focused more on intracellular and extracellular biochemical signaling. For example, Jilkine et al investigated multiple possible interaction schemes between the Cdc42, Rac, and Rho GTPase proteins that give rise to physiological spatial polarization (28). Mare et al extended a spatially distributed model of Rho GTPase proteins interacting with actin filament dynamics, coupled to a Cellular Potts model accounting for spatial aspects of cellular polarization and protrusion (27). Merks and colleagues have developed several Cellular Potts modeling approaches, coupling cell-cell and cell-extracellular matrix interaction energetic constraints with extracellular biochemical concentrations, to study collective cell migration and tissue dynamics in several physiological settings, including angiogenesis and tumor invasion (35–38).

Thus, many of these prior studies have accounted for detailed representations of either mechanical or biochemical signaling. Our approach extended from prior work from Zmurchok et al (19) represented a compromise between accounting for both mechanical and biochemical signaling and feedback, while still utilized a minimal approach that facilitated wide-ranging parameter studies to characterize model behavior. In comparing our work with this prior study, Zmurchok and colleagues also observed center-to-periphery Rho GTPase propagating waves in one-dimensional simulations of stationary tissue (comparable to Fig. 5A), suggesting that junctional forces are not necessary for this mechanochemical synchronization (since junctional forces are not specifically represented in Zmurchok et al). Interestingly, by accounting for individual cell front-back polarity, we find that the mechanochemical waves also reorient to a tissue back-to-front and front-to-back propagating pattern in a migrating tissue (Fig. 5B).

### Physiological significance

Despite the minimal nature of our model, we can gain useful insights for understanding fundamental *in vivo* processes involving cell migration, in particular to understand the role of forces in subsets of collective cell migration. For example, chain migration of cells is defined as cells that start as a cluster and delineate away in a single file line toward a chemotactic gradient (39) and has been demonstrated *in vivo*. In the xenopus embryo neural crest, cell migration depends upon non-canonical Wnt activation of RhoA until they form cell-cell contacts with another cell, causing them to change direction (40). This study demonstrated the concept of cell-cell contacts influencing migration speed and direction, but did not take into account the force at these contacts nor the traction force of the system. The chain migration of neural crest cells has been further probed *in vitro*, showing that high cell tension at the periphery inhibits spreading and migration, while lower tension in the leader cell promotes leader cell spreading and higher traction forces at later stages of migration (41).

Leader cells have also been shown to be important in cancer metastasis, driving the collective migration of multi-cellular groups of cancerous cells into the surrounding tissue (42). Interestingly, simulations of heterogeneous tissues predict that a single migratory leading cell can sufficiently drive migration of a tissue comprised of stationary trailing cells, suggesting that metastasis may not require a population of primarily motile cancerous cells. Our model may also be compared with physiological branching morphogenesis. In branching morphogenesis occurring in the lung, kidney, vascular system, prostate, mammary gland, and kidney to form tubular branching structures, a leader or “tip” cell responds to a chemotactic or durotactic gradient and protrudes while bringing along follower or “stalk” cells through adhesion junctions. Several studies have shown that mechanically sensitive molecules such as YAP (43), integrin *β*1 (44), and the TRPV4 ion channel (45), are needed for proper branching morphogenesis, yet measurement of cell-cell force during this process *in vivo* is limited. Since *in vivo* collective cell migration is difficult to quantify, especially with respect to cell-cell and cell-matrix forces, our model provides a method for understanding this fundamental aspect of development and disease.

Perhaps most relevant to this model is experimental work by Fredberg and colleagues in which measurement of traction forces of migrating sheets of epithelial cells showed significant mechanical forces not only at the leading edge but several rows behind (46). However, in contrast with most of our model predictions (e.g., Figs. 7 and 8), the traction force measurements of Fredberg and colleagues demonstrate highest cell-cell junction forces for cells far away from the leading edge. While there are many physical differences between the two-dimensional epithelial sheet migration and one-dimensional migration studied here, these differences could alter tension-feedback in subtle ways that result in junctional force spatial patterns comparable to Figure 9A, in which larger junctional forces are present away from the leading edge.

An interesting extension of our current work would be to investigate collective cell migration in the context of cell jamming, in which key cellular and tissue properties, such as motility, density, size, stiffness, cell-cell adhesion, etc., are considered as different “dimensions”, and transitions along these dimensions out of a “jammed” or stationary state can result in different forms of collective migration (47). Our model framework is ideal to consider these issues, e.g. by incorporating mechanochemical regulation of cellular proliferation and cell size, while still maintaining a minimal representation that can be extensively explored in parameter space.

### Limitations

The model formulation used in this study is inherently minimal by design. specifically, we did not consider interactions between different Rho GTPase proteins, namely the crosstalk and mutual feedback between Cdc42, Rho, and Rac, nor the well-established subcellular front-back spatial localization of these proteins during polarization and migration (17, 48). Prior work using detailed multiscale models from the Edelstein-Keshet group has shown that spatial segregation of the Rho GTPase proteins is critical for robust polarization (11, 27, 28). Similarly, while cellular extension, contraction, and migration are inherently a two- or three-dimensional behavior, as focal adhesion formation and integrin binding occurring over a distributed subcellular volume and surface area, respectively, our model minimally represents a single direction or dimension for migration. It would be of interest to consider more spatially detailed representations of these processes; however it may be computationally prohibitive or at a minimum expensive to account for this level of biophysical detail (i.e., accounting for subcellular spatial concentration distributions) in a multicellular tissue. Finally, our model does not consider feedback between junctional forces and intracellular cytoskeletal signaling, such as global rearrangements of the cytoskeleton, which in turn may influence properties of Rho GTPase signaling and/or mechanical properties related to cell contractility or focal adhesion formation. Incorporating these relationships is a natural extension of our work presented here and is an area of interest that we will consider in future work. Our initial focus was to investigate fundamental relationships between key processes governing collective cell migration, specifically intracellular Rho GTPase activity, cellular mechanical forces, and intercellular junctional force interactions.

## CONCLUSION

In this study, model predictions illustrate that collective cell migration is governed by emergent mechanochemical interactions, propagating waves of Rho GTPase activity that synchronize mechanical contraction and extension throughout the tissue. Cell-cell junctional forces exhibit a distinct spatial pattern, with larger forces near the leading edge of the tissue and larger junctional forces associated with faster collective migration. Finally, simulations in both homogeneous and heterogeneous tissue illustrate that collective migration does not depend simply on the velocity of individual cells comprising the tissue, but additionally on the mechanochemical interactions that govern intercellular junctional forces and synchronization within the tissue.

## AUTHOR CONTRIBUTIONS

SHW designed the research. JB and SHW carried out all simulations and analyzed the data. DEC, RLH, and SHW wrote the article.

## ACKNOWLEDGMENTS

This work was supported by the National Institutes of Health via award R01GM122855 (SHW) and R35GM119617 (DEC).

