## Supplemental Material for "Mechanochemical coupling and junctional forces during collective cell migration"

### SUPPLEMENTAL FIGURES

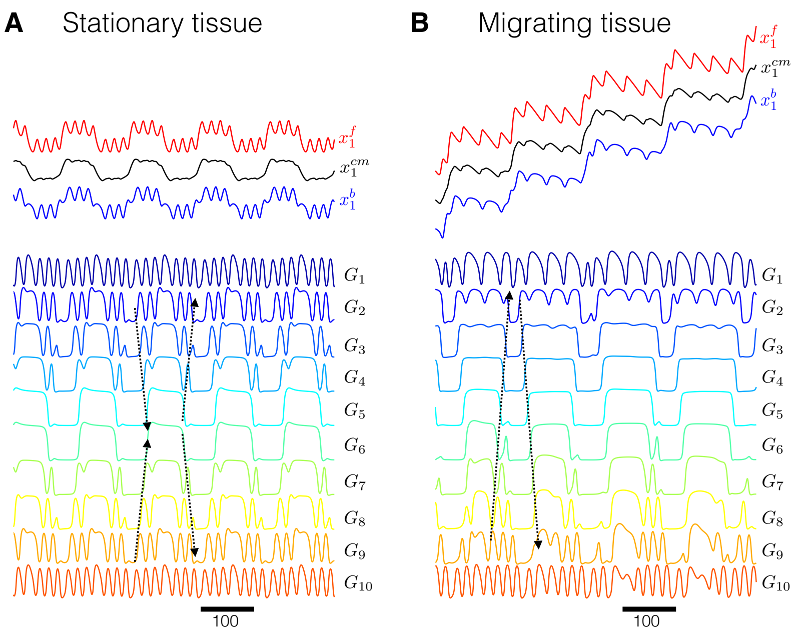


Figure S1: Dynamics of Rho GTPase activity in homogeneous tissue. (Top) The time course for cell positions for the front cell in the tissue (front (red) $x_1^f$, center (black) $x_1^{cm}$, and back (blue) $x_1^b$), and (Bottom) Rho GTPase activity ($G_i$, for $i = 1, 2, \ldots, n = 10$) are shown in (A) stationary and (B) migrating tissue. Dashed arrows highlight propagating waves of high and low Rho GTPase activity. Parameters: $\beta = 0.16$, (A) $\delta = 0$, (B) $\delta = 0.9$.

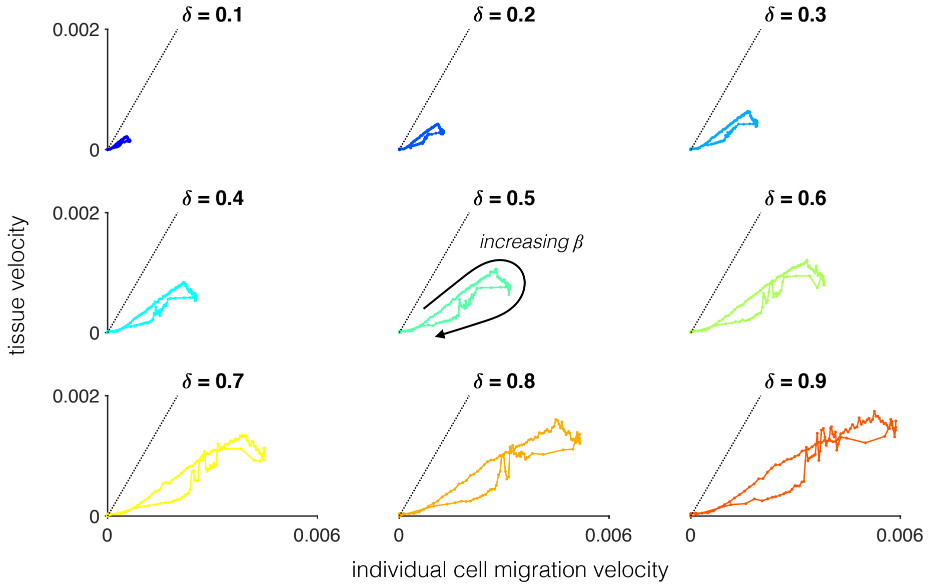


Figure S2: Proportionality of homogeneous tissue velocity and individual cell migration velocity. For a given value of front-back polarity parameter $\delta$, the tissue velocity is measured and plotted against the corresponding individual cell migration velocity, as tension-feedback parameter $\beta$ is varied. Black dashed line is a line with slope 1. For low $\delta$ and low $\beta$ values, individual and tissue migration velocities are proportional. However, differences in the optimal $\beta$ value between individual and tissue migration result in a parameter regime for which tissue velocity decreases for larger individual cell migration velocity. Parameters: Tissue size of 10 cells.

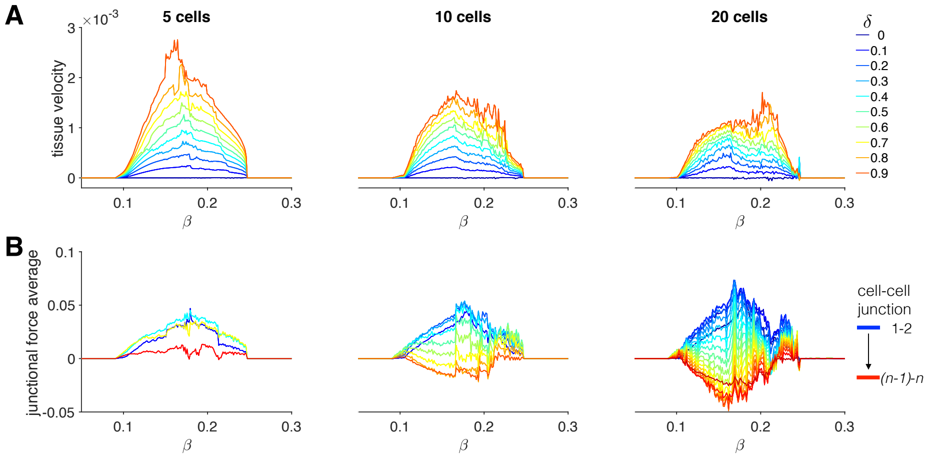


Figure S3: Collective cell migration properties depend tissue size. (A) Tissue velocity is shown as a function of $\beta$ and different values of $\delta$, for tissue sizes of 5, 10, and 20 cells. Dependence on $\beta$ and $\beta$ are similar for different tissue sizes; however, velocity in general decreases as tissue size increases. (B) Junctional force averages from the junctions at the tissue front (blue) to the tissue back (red) are shown as a function of $\beta$, for tissue sizes of 5, 10, and 20 cells. Parameters: (B) $\delta = 0.5$

### SUPPLEMENTAL TABLES

Table S1: Model parameters for individual and collective cell migration.

| Parameter | Value(s) |
|---|---|
| $b$ | 0.1 |
| $\gamma$ | 1.5 |
| $G_T$ | 2 |
| $l_0$ | 1 |
| $G_h$ | 0.3 |
| $p$ | 4 |
| $q$ | 4 |
| $\phi$ | 0.75 |
| $\alpha$ | 10 |
| $\beta$ | [0.05, 0.30] |
| $\epsilon^b$ | 0.1 |
| $\delta$ | [0,1) |
| $k$ | 1 |
| $\kappa_{junc}$ | 14 |
| $L_{junc}$ | 0.05 |

### SUPPLEMENTAL MOVIES

Movie S1: Migration of an individual cell. (Top) The position of the cell center of mass $x^{cm}$ and Rho GTPase activity $G$ are shown as a function of time. Scale bar = 50. (Middle) Representation of the cell position are shown for different times. Color of the cell corresponds to Rho GTPase activity, with blue denote a low level (0.2) and red denotes a high level (0.5). (Bottom) Position of the cell front (red), center (black), and back (blue) are shown as a function of time. Note time is on the y-axis. Scale bar for middle and bottom panels = 0.5. Movie corresponds with results shown in Fig. 3B. Parameters: $\beta = 0.16$, $\delta = 0.5$.

Movie S2: Migration for individual cells with increasing front-back polarity. Representation of cell position for individual cells with different values of front-back polarity parameter $\delta$ are shown for different times. Note that the color corresponds to the $\delta$ value. Scale bar = 0.5. Parameters: $\beta = 0.16$.

Movie S3: Dynamics of a stationary tissue. (Top) Representation of cell positions in a stationary tissue. Color of the cell corresponds to Rho GTPase activity, with blue denote a low level (0.2) and red denotes a high level (0.53). (Bottom) Position of cell fronts (red), centers (black), and backs (blue) are shown as a function of time. Note time is on the y-axis. Scale bar = 0.5. Movie corresponds with results shown in Fig. 4 and 5A. Parameters: $\beta = 0.16$, $\delta = 0$.

Movie S4: Dynamics of a migrating tissue. (Top) Representation of cell positions in a migrating tissue. Color of the cell corresponds to Rho GTPase activity, with blue denote a low level (0.2) and red denotes a high level (0.53). (Bottom) Position of cell fronts (red), centers (black), and backs (blue) are shown as a function of time. Note time is on the y-axis. Scale bar = 0.5. Movie corresponds with results shown in Fig. 5B and 6C. Parameters: $\beta = 0.16$, $\delta = 0.9$.

Movie S5: Mechanochemical resonance during collective cell migration. Representation of cell positions in migrating tissues with small differences in tension feedback parameter $\beta$. Color of the cell corresponds to Rho GTPase activity, with blue denote a low level (0.2) and red denotes a high level (1.1). Scale bar = 0.5. Movie corresponds with results shown in Fig. 9. Parameters: $\delta = 0.9$.
